# Spatial Dynamics and Functional Connectivity in EEG: Insights from Lexical Processing

**DOI:** 10.1101/2025.04.29.651245

**Authors:** Omar Aguilar, Ramon Grullon

## Abstract

Lexical processing is a core cognitive function involving the integration of semantic and symbolic information across distributed brain networks. In this study, we investigate the spatial dynamics and functional connectivity underlying lexical categorization using electroencephalography (EEG) data from a silent reading task. Participants were presented with words from two categories—social and numeric—while EEG signals were recorded and analyzed. Frequency band power and inter-electrode coherence were extracted as features to train Random Forest classifiers for word category prediction. The spatial dynamics model, based on band power from selected electrodes, achieved 100% classification accuracy and identified region- and frequency-specific contributions, notably in the theta and alpha bands across parietal and occipital regions. A complementary functional connectivity model achieved 85% accuracy, highlighting the role of inter-regional coherence in lexical differentiation. These findings demonstrate the potential of combining traditional EEG analysis with machine learning for decoding semantic categories, providing foundational insights for future brain-computer interface applications.

## I. Introduction

Understanding the spatial dynamics and functional connectivity of the brain during lexical processing is a crucial step toward developing advanced brain-computer interfaces (BCIs). These interfaces aim to decode and classify brain signals, potentially allowing individuals with motor impairments to communicate more effectively [1]. By identifying specific spatial and connectivity patterns in the brain, researchers can gain deeper insights into how different regions coordinate to process linguistic information.

Lexical processing is a fundamental cognitive task involving the interpretation and integration of semantic and symbolic information. It is well-known that distinct frequency bands in EEG data correspond to specific brain states, ranging from low-frequency delta waves to high-frequency gamma waves. However, the relative importance of these frequency bands in distinguishing different types of words remains an open question in the field. Moreover, the functional connectivity between brain regions, characterized by the synchronization of neural signals, plays a vital role in information integration, yet its contribution to lexical processing classification is not fully understood.

This study investigates these phenomena by analyzing EEG data collected from participants performing a controlled lexical task. The experiment involved four participants silently reading words from two categories, social and numeric, while their neural activity was recorded. With 320 trials per participant, the task allowed for robust data collection and analysis [2]. Using advanced machine learning methods, such as Random Forest models, and neurophysiological techniques, such as coherence analysis, this research aims to uncover key spatial patterns and connectivity dynamics that underpin lexical processing.

By addressing these questions, this work provides critical insights into the neural mechanisms of language comprehension. It also highlights the potential for developing BCIs capable of distinguishing cognitive tasks based on brain signals. these findings may pave the way for improved communication technologies for individuals with severe motor impairments, as well as a deeper understanding of how the brain processes different categories of information.

## II. Literature Review

The analysis of neural and behavioral data has long leveraged traditional methods such as band power and coherence analysis to uncover insights into brain function. Band power, derived from EEG frequency domain representations, has been a central feature in understanding spatial dynamics across specific brain regions. For instance, studies on lexical processing tasks have identified significant changes in EEG band power, particularly within alpha (8–12 Hz) and beta (13–30 Hz) frequency bands. Decreased alpha power and increased beta power in temporal and frontal cortices have been consistently associated with semantic and phonological processing demands [6]. These findings underscore the importance of band power in the decoding of cognitive and linguistic processes.

Coherence analysis complements these findings by examining the synchrony between EEG signals across different brain regions, providing a window into functional connectivity. Research has highlighted the role of coherence, particularly in conditions such as Alzheimer’s disease, where decreased interhemispheric alpha-band coherence reflects disrupted brain network dynamics [11]. In behavioral tasks, coherence analysis has demonstrated how different brain areas interact, emphasizing the importance of network-level connectivity over isolated regional activity [9].

Despite the robustness of these traditional methods, their utility has often been limited to correlation-based analyses, which are constrained by linearity assumptions and fail to capture complex interdependencies. Machine learning offers an avenue to address these limitations by enabling the classification of cognitive states and identifying critical neural features [10]. Recent advances have utilized models such as support vector machines (SVMs) and random forests to quantify the importance of specific EEG electrodes in distinguishing task conditions. By integrating nonlinear relationships, machine learning approaches can provide richer insights into neural dynamics, surpassing the capabilities of traditional correlation methods [4].

However, notable gaps in the literature persist. First, while band power and coherence are widely used, their integration into machine learning frameworks for classification remains underexplored. This project seeks to bridge this gap by combining band power and coherence features to improve model interpretability and performance. Second, the specific application of these techniques to lexical processing tasks is limited. By focusing on this domain, this work aims to elucidate how EEG features contribute to understanding language processing mechanisms. Lastly, coherence analysis is rarely incorporated into machine learning workflows, despite its potential to enhance classification accuracy and reveal inter-regional dynamics [7].

This project addresses these gaps by advancing the field of neural and behavioral data analysis. Through the innovative combination of traditional EEG analysis methods and machine learning, it seeks to provide a comprehensive framework for understanding brain function and its relationship to behavior. Such an approach not only enhances predictive accuracy but also deepens insights into the intricate interplay of neural dynamics, setting a foundation for future research in cognitive neuroscience.

## III. Methods

### Experimental Design

The data used in this analysis is from a paper published by researchers at the Lulea Institute of Technology [2]. This dataset does involve functional magnetic resonance imaging (fMRI) data, but for the purpose of this study, only the EEG data was used. The experiment was conducted with five participants who completed a controlled lexical processing task, but one patient was excluded due to high fluctuations during the EEG recording. In the specific application of this study, only data from sub-02 was used, as it yielded the most interpretable and clear results. Participants were each presented with one of eight words displayed on a screen, categorized into two groups: four social words and four numeric words. The social words include *child, daughter, father, and wife*, while the number category contains *four, three, ten, and six*. Participants were instructed to silently read the word on the screen and repeat it internally. Each trial lasted 4 seconds, comprising 1 second of fixation, 2 seconds of inner-speech processing, and 1 second of rest. In total, 320 trials were conducted per participant, ensuring sufficient data for robust analysis.

### EEG Preprocessing and Feature Extraction

Electroencephalogram (EEG) data were recorded from multiple electrodes placed on the scalp using the international 10-20 system. The raw EEG signals were preprocessed using MEG + EEG Analysis & Visualization (MNE) python library to remove artifacts and baseline drifts. Electrode channels were assigned according to the montage. A bandpass filter was applied to the signal from 1 to 50 Hz using a windowed time-domain design (firwin) method and a phase shift of zero. The reference channel was set to the “Cz” electrode and averaged. Independent component analysis (ICA) was performed, which included the exclusion of electrooculogram (EOG) channels. The signal for each electrode was parsed according to the experimental paradigm for each trial. From this, the 320 trials were created with lists of the time points for each given electrode.

From the original set of 64 electrodes, only 25 were used from the regions of interest for a lexical processing task (Table I. Frequency domain analysis was performed to extract power in specific frequency bands (II), providing a metric indicative of neural activity within those bands.

**TABLE I:** Selected electrodes - This chart includes the brain regions shown to be related to lexical processing along with the selected electrodes from those regions.

| Brain Region | Electrodes |
| --- | --- |
| Occipital | O1, O2, Oz |
| Parieto-Occipital | POz, PO3, PO4, PO7, PO8 |
| Frontal | F3, F4, Fz, F7, F8 |
| Anterior Frontal | AF3, AF4, AF7, AF8, Fpz |
| Temporal | T7, T8 |
| Central | Cz, FCz |
| Parietal | Pz, CP1, CP2 |

**TABLE II:**
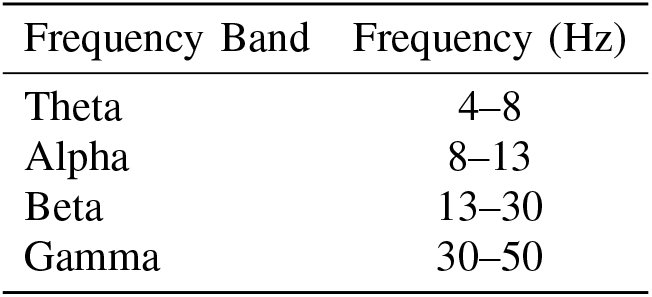
Frequency band chart - These are the frequency bands that were observed, along with their respective frequency values.

To compute the power within these frequency bands, Welch’s power spectral density (PSD) was calculated using the python scipy library. In preparation for the connectivity analysis, electrode pairs were made. Bandpass filters were applied to the electrode pair signals within the predefined frequency ranges, and coherence was calculated using the between the filtered signals for each frequency band.

### Model Training, Optimization, & Evaluation

For this study, Random Forest Classification was used to determine the category of the word (social vs. numeric) in a specified trial. Random Forest is an ensemble learning method composed of multiple decision trees (a.k.a. estimators), which improves accuracy and reduces overfitting [3]. Each decision tree is trained on a random subset of the data and a random subset of features, creating diverse trees that capture different patterns. The final classification decision is determined by a majority vote across all trees.

One of the main reasons this model was chosen is its *feature importance* attribute, which quantifies how much each input feature contributes to the model’s predictive performance. Feature importance is typically derived from the *impurity reduction* achieved when a feature is used to split a node in a decision tree. For classification tasks, impurity is often measured using the Gini index:

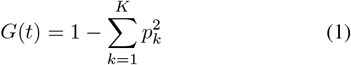

where *G*(*t*) is the Gini impurity at node *t, p*_*k*_ is the proportion of samples belonging to class *k*, and *K* is the total number of classes.

When a feature is used to split a node, the *decrease in impurity* (information gain) is computed as:

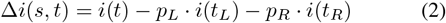

where *s* is the split criterion (feature and threshold), *i*(*t*) is the impurity at the parent node, *i*(*t*_*L*_) and *i*(*t*_*R*_) are the impurities of the left and right child nodes, and *p*_*L*_, *p*_*R*_ are the proportions of samples that go to each child.

The feature importance for feature *f* in a single decision tree is then calculated by summing the impurity decreases at all nodes where *f* is used:

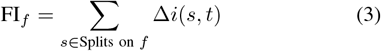

For the entire Random Forest, the overall feature importance is obtained by averaging over all *T* trees:

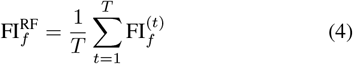

This framework not only provides robust classification accuracy, but also yields interpretable insights into which features (such as frequency band power or coherence values) are most critical in distinguishing lexical categories.

Two random forest models were made in this study, the first being a model that focuses on spatial dynamics within the different brain regions. This model was trained on frequency band powers for each electrode and trial. From this, the importance for each feature was used to determine which regions of the brain contributed the most to classification between social and numeric words based on the location of the electrode. Also from this model, the average feature importance of each frequency band, regardless of electrode location, was examined to see which frequency bands played the largest part in classification.

The second classifier was made to focus on functional connectivity between electrodes within the brain. This model was trained on coherence values of bandpass filtered electrode pair signals within different frequency bands. From this, feature importances were extracted to examine the most important connections within this classification task.

For both of these models, hyperparameter optimization was performed for the Random Forest Classifiers using a Grid Search Cross-Validation algorithm. In this, different values or settings are listed. The different combinations of parameters are trained across five different folds to determine the best set of parameters for each model. This cross-validation method prevents overfitting by training across different folds while still selecting the best set of parameters using accuracy. After selecting the optimal hyperparameters, the models were evaluated using standard accuracy and confusion matrices for a more detailed representation of classifier performance.

## IV. Results

### Spatial Dynamics Model

This Random Forest Classifier achieved an accuracy of 100% in the classification of social vs. numeric words. From this model, a topographical scalp map (Figure 3) was generated to visualize which brain regions contributed the most to classification based on frequency band power in specifc electrodes.

**Fig. 1:**
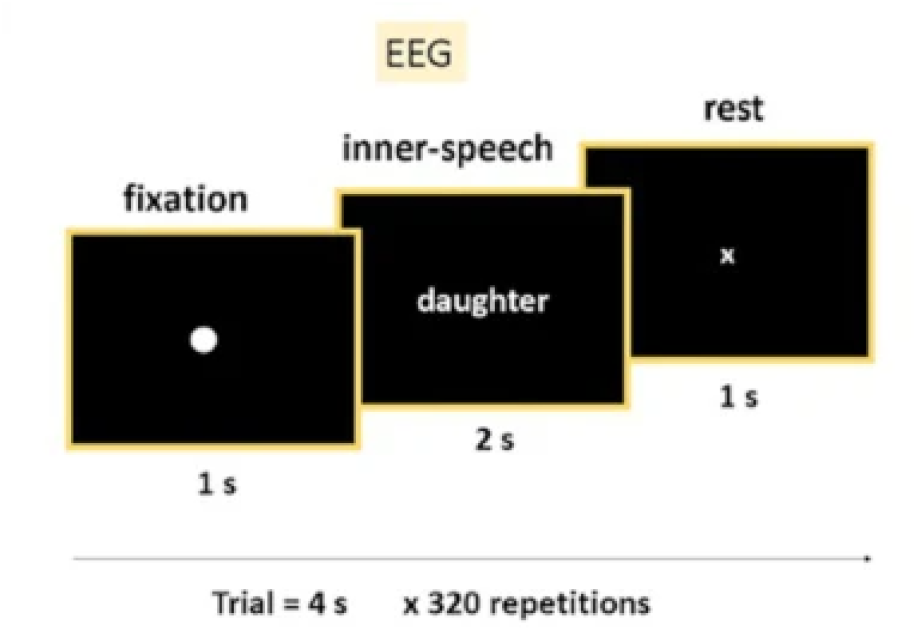
Experimental paradigm for EEG testing - Each trial consists of one second fixation, two seconds inner-speech, and one second rest. This process is repeated 320 times, showing a random word each time [2].

**Fig. 2:**
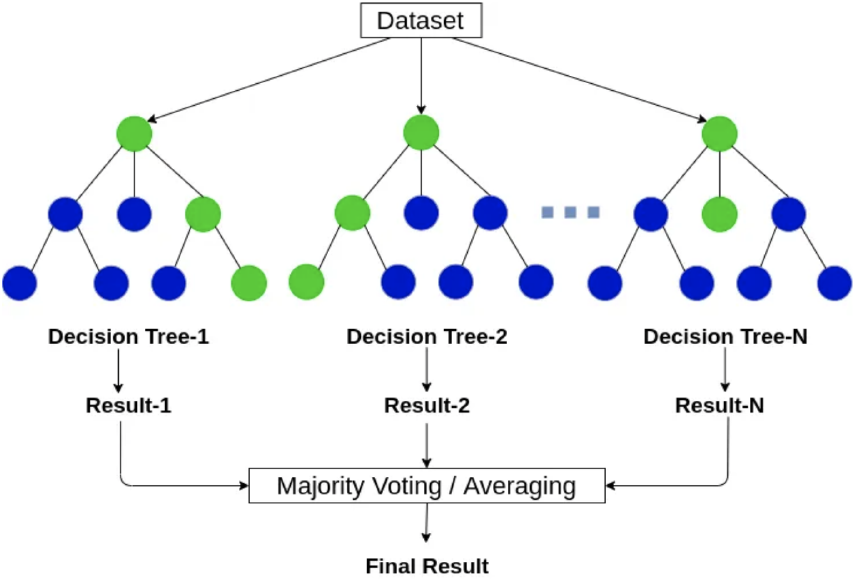
Random forest classification structure, portraying how a classification decision is made. [13]

**Fig. 3:**
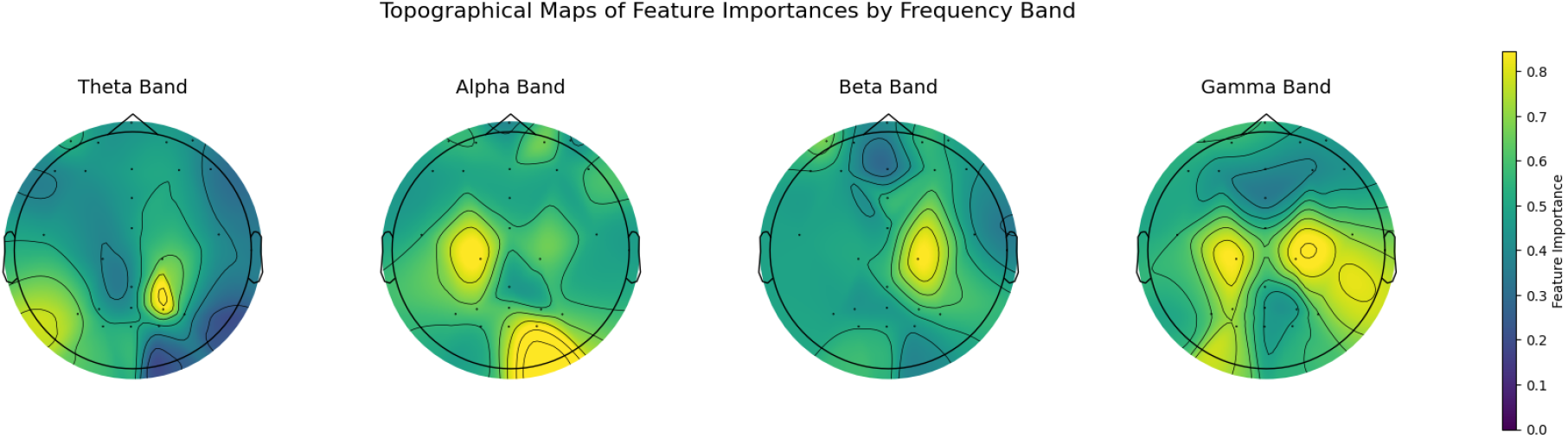
Topographical maps of feature importances by frequency band - This figure gives a visualization to feature importance based on frequency band power for specified electrodes. The location of intensity is based upon the electrode placement within the montage.

High feature importance in the right parietal region of the theta band can be seen from the figure. This suggests involvement in semantic integration and symbolic processing, leading to distinct cognitive processes for social and numeric words in this task. Dominance in the left central-parietal and occipital regions of the alpha band is also apparent. This outcome suggests that analytical vs. semantic processing was a differentiating factor in classification. There isn’t much in the beta band, but high importance is seen to be concentrated in the right central parietal region. This reflects cognitive and motor demands such as focused attention and cross-modal integration play a key role in the classification task. Lastly, high feature importance across the central parietal, temporal, and occipital regions are seen in the gamma band. This points to higher-order cognitive processing, visual attention, and semantic integration playing crucial roles differentiating between social and numeric words.

Figure 4 was also generated to show the frequency band power feature importance. In this, the theta and beta bands exhibited the highest average feature importance based on frequency band power. This suggests that the task heavily engages in semantic processing (theta) and focused attention (beta), which are essential for distinguishing between social and numeric words.

**Fig. 4:**
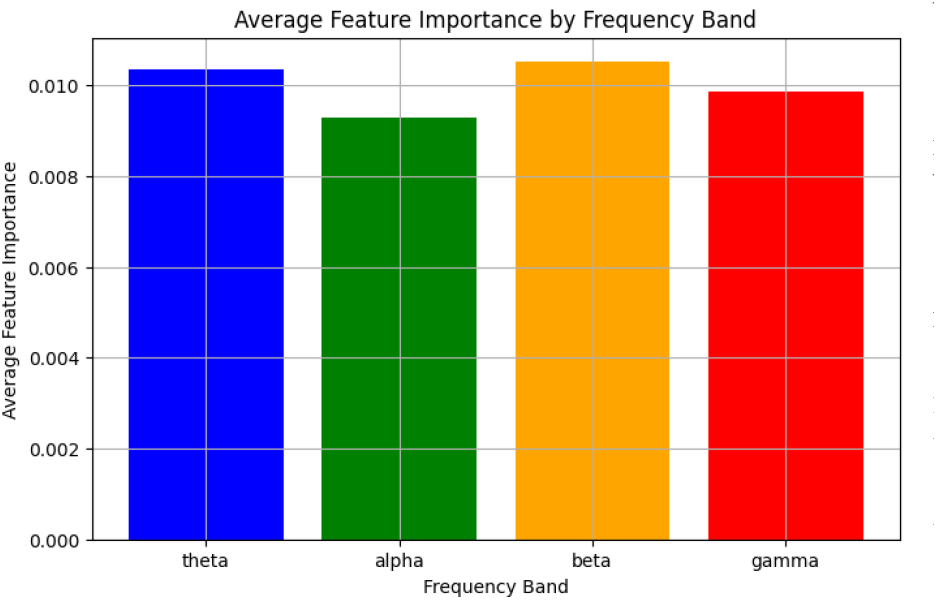
Average feature importance by frequency band - This figure highlights the importance of each frequency band in the classification task.

**Fig. 5:**
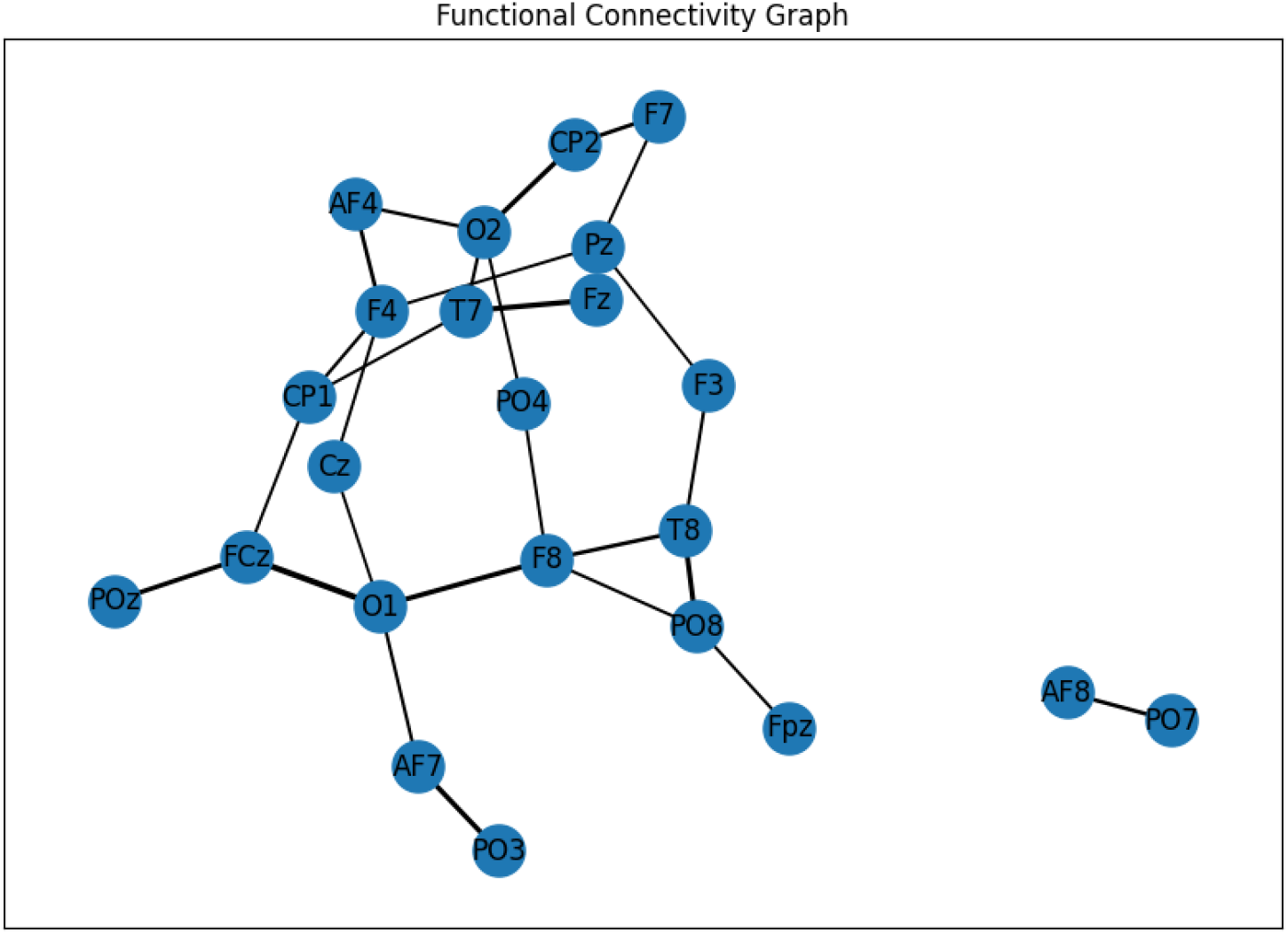
Functional connectivity graph - This figure shows the strength of coherence between signals in electrode pairs, along with the feature importance of the electrode pairs.

### Functional Connectivity Model

In this model, an accuracy of 85% was achieved. From this, a functional connectivity graph was generated using a spring layout. The node proximity represents the strength of the coherence between an electrode pair. The line thickness corresponds to the feature importance, therefore, the most important connections in the classification task will have the thicker lines. To generate this graph, a threshold was set for feature importance to only show the top percentage of connections. Because of this, isolated groups can be formed. No connection between nodes indicates it did not meet the threshold of importance.

In this figure, electrodes from the central and parietal regions are densely clustered. This indicates strong coherence in these areas and suggests that these regions may be crucial for information integration during the task. The thicker lines connecting electrodes like O1-FCz and O2-CP2 indicate that the connectivity between these regions is especially significant for the model’s classification. This implies that these connections play a key role in distinguishing between conditions in the task. The isolated connection between AF8 and PO7 reveals an important finding. While this connection is important enough to be on the graph, it is not heavily integrated with the primary network. This potentially reflects a form of specialized processing that is rendered important in differentiating between social and numeric words.

## IV. Discussion

The results of our analysis provide meaningful insights into the neural mechanisms underlying lexical processing, specifically as they pertain to the differentiation between social and numeric words. The spatial dynamics model, which achieved perfect classification accuracy, highlighted the role of various brain regions in lexical categorization. The high feature importance observed in the right parietal region within the theta band indicates that this area is involved in semantic integration and symbolic processing. This aligns with existing research that associates theta band activity with language comprehension and cognitive load during complex processing tasks. Additionally, the prominence of the left central-parietal and occipital regions in the alpha band suggests that distinct patterns of analytical versus semantic processing play an important role in discriminating between the two categories of words.

These findings imply that the differentiation between social and numeric words relies heavily on both localized and distributed neural activity. The machine learning models reveal an underlying structure in the neural data that is distinctly associated with semantic processing, emphasizing the importance of specific frequency bands and their regional significance. The spatial dynamics model, with its perfect accuracy, suggests that certain brain regions exhibit highly distinctive activation patterns for social versus numeric words, allowing for clear classification. This supports the hypothesis that lexical processing is not just localized but also involves different neural frequencies that encode unique aspects of semantic understanding.

When compared to existing literature, our findings are consistent with the role of the theta and alpha bands in language comprehension and semantic processing. Previous studies have established that theta band activity is linked to memory retrieval and semantic integration, which aligns with the observed feature importance in the right parietal region [6]. The dominance of the alpha band in the left central-parietal and occipital regions also supports prior research suggesting that alpha oscillations are crucial for attentional processes and semantic suppression [5]. However, the relatively lower importance of the beta band contrasts with some literature that associates beta activity with language and motor functions, suggesting that these roles may be less pronounced in silent reading tasks as used in our study [12].

The functional connectivity model, which attained 85% accuracy, further emphasizes the importance of coherence between specific brain regions in distinguishing lexical categories. The dense clustering of electrodes in the central and parietal regions suggests that these areas are critical for integrating information during the task. The strong connections between electrodes such as O1-FCz and O2-CP2 point to significant inter-regional communication that supports differentiation between social and numeric words. Interestingly, the isolated connection between AF8 and PO7 may indicate a specialized form of processing that, while not central to the main network, still plays a crucial role in this classification task. This finding could indicate a unique pathway for processing specific types of semantic information, emphasizing the complexity of lexical processing that involves both central and specialized networks.

Despite the promising findings, there are several limitations to this study that should be acknowledged. The perfect classification accuracy of the spatial dynamics model may suggest potential overfitting, especially given the relatively small dataset used (data from only one participant). This raises questions about the generalizability of the model to other individuals or broader populations. The functional connectivity model, while insightful, only achieved 85% accuracy, indicating that the coherence-based features may not fully capture the complexity of lexical processing or that there may be other factors influencing connectivity patterns that were not accounted for. Further research with larger participant samples and cross-validation is needed to validate these findings and improve the robustness of the models.

Potential sources of error also include the preprocessing steps, such as Independent Component Analysis (ICA) and the choice of frequency bands. While these steps are necessary to remove noise and artifacts, they also have the potential to inadvertently remove meaningful signals. Moreover, the use of a fixed reference electrode (Cz) might have influenced the observed connectivity patterns, potentially biasing the results towards central regions. Future studies could benefit from exploring alternative referencing methods and including additional participants to enhance the generalizability of the results.

Another limitation in this study involves a possible language barrier that could pose issues in the lexical processing task. This experiment was performed at Lulea University of Technology in Sweden, and each participant spoke a different native language. The native language of sub-02 was Bengali. Though the words still might have the same meaning, it might elicit different activation patterns from words seen in the subject’s native tongue. Future studies could explore the differences in words shown in a subject’s native language vs. a learned language. Future research could also inspect emotional connections to some of the social words through a pre-experimental survey, which may give more insight to specific activations patterns in a participant.

Together, these findings reinforce the notion that lexical processing involves complex interactions between multiple regions and frequencies, reflecting both local activity and global network connectivity to accomplish the task at hand. The models provide valuable insights into the neural underpinnings of lexical categorization, but also highlight the need for careful consideration of model limitations and data constraints to ensure reliable interpretations.

Beyond providing insight into the neural mechanisms of lexical processing, these findings have important implications for brain-computer interface (BCI) development. By identifying distinct spatial and functional connectivity patterns associated with semantic categories, this study lays the groundwork for EEG-based cognitive decoding systems. In particular, the clear regional and frequency-specific signatures revealed by the spatial dynamics model suggest that compact, targeted EEG montages could be designed to classify cognitive states with minimal electrode coverage, improving portability and user comfort in real-world BCIs. Furthermore, the functional connectivity model highlights the potential for dynamic, network-based features to enhance decoding accuracy in noisy or mobile environments. Future research could extend this framework to online decoding scenarios, enabling real-time cognitive state monitoring and augmentative communication tools for individuals with impaired motor function.

## VI. Conclusion

In conclusion, this study demonstrates the power of machine learning, particularly Random Forest models, in uncovering patterns in complex neural data related to lexical processing. The spatial dynamics model revealed distinct regional and frequency-specific activation patterns that differentiate between social and numeric words, while the functional connectivity model highlighted the significance of inter-regional coherence in lexical categorization. These findings underscore the value of machine learning in extracting meaningful information from intricate biological signals, offering a deeper understanding of how different brain regions contribute to language processing. Future research should focus on expanding the participant pool to validate these findings across a more diverse population, addressing potential biases introduced by the current dataset, and refining the preprocessing steps to minimize the risk of losing important signals. Additionally, exploring alternative models and more advanced connectivity analyses may further elucidate the underlying neural mechanisms of lexical processing, potentially paving the way for improved interpretability and practical application in cognitive neuroscience.

